# Systematic Dissection of Sequence Elements Controlling σ70 Promoters Using a Genomically-Encoded Multiplexed Reporter Assay in *E. coli*

**DOI:** 10.1101/207332

**Authors:** Guillaume Urtecho, Arielle D. Tripp, Kimberly D. Insigne, Hwangbeom Kim, Sriram Kosuri

## Abstract

Promoters are the key drivers of gene expression and are largely responsible for the regulation of cellular responses to time and environment. In *E. coli,* decades of studies have revealed most, if not all, of the sequence elements necessary to encode promoter function. Despite our knowledge of these motifs, it is still not possible to predict the strength and regulation of a promoter from primary sequence alone. Here we develop a novel multiplexed assay to study promoter function in *E. coli* by building a site-specific genomic recombination-mediated cassette exchange (RMCE) system that allows for the facile construction and testing of large libraries of genetic designs integrated into precise genomic locations. We build and test a library of 10,898 σ70 promoter variants consisting of all combinations of a set of eight ‐35 elements, eight ‐10 elements, three UP elements, eight spacers, and eight backgrounds. We find that the ‐35 and ‐10 sequence elements can explain approximately 74% of the variance in promoter strength within our dataset using a simple log-linear statistical model. Simple neural network models explain greater than 95% of the variance in our dataset by capturing nonlinear interactions with the spacer, background, and UP elements.

## Introduction

Promoters are the key regulators of gene expression and largely control developmental and environmental responses in all living organisms^1-3^. Decades of studies on bacterial and phage promoters have elucidated many of the essential proteins and basic sequence motifs necessary for initiating transcription. In *E. coli*, transcription requires the polymerase holoenzyme which consists of a core set of five subunits, as well as one of seven known sigma factors^4-6^. Sigma factors provide much of the sequence specificity for bacterial promoters, and the prevalence of each factor varies based on environmental conditions^7,8^. Under standard growth conditions, most active promoters are transcribed by σ70^5^. σ70-dependent promoters are composed of discrete sequence elements that cooperatively determine expression, including two conserved hexamers centered ‐10 and ‐35 bases upstream of the transcription start site that directly interact with the σ70 subunit^9^. Other sequence elements are known to be important including the nucleotide content and length of the spacer region between the ‐10 and ‐35^10^, an UP element upstream of the ‐35 that can anchor the RNA polymerase (RNAP) α-subunit to enhance promoter recognition^11^, and the local sequence context surrounding these elements^12^.

Despite the apparent simplicity of the process and decades of genetic and biochemical dissection, we still lack answers to basic questions surrounding bacterial transcription. For example, given an arbitrary sequence, we largely do not have the ability to know (1) if it is a promoter, (2) its strength, and (3) its regulation. Thus far most approaches try to understand the relationships between sequence elements that comprise the full promoter using reverse genetic approaches where they characterize multiple variants of an element in a single promoter context^10,13^. Although these studies have revealed the contributions of individual sequence elements, the effects of these variants are often inconsistent between promoters, conceivably due to higher order relationships between sequence elements^14^. Conceptually, a simple way to tease apart these relationships is to test a wide variety of element combinations across a variety of backgrounds. However, increasing the number of element combinations and backgrounds quickly surpasses the number of constructs that can be practically tested using traditional means. Developing novel approaches to test vastly more designs will help us understand the behavior of sequence elements across different contexts, and more broadly allow us to explore relationships between promoter sequence and function.

Massively Parallel Reporter Assays (MPRAs) are a new class of experiments that test large numbers of designed genetic variants for functional activity^15,16^. We and others have used MPRAs to quantify expression of large promoter or enhancer libraries for both transcriptional and translational activities^17-27^. However, there are several limitations when using these systems to study promoter function in bacterial systems. First, since many current systems rely on flow cytometry and sorting, reporter expression levels must be relatively high to detect signal, limiting the quantitative range of promoter studies. Second, these systems often measure protein production rather than RNA, making it difficult to decouple transcriptional and translational processes. Third, although there are many bacterial MPRAs that measure transcriptional readout directly, most systems accomplish this by quantifying unique sequence tags located on the 5’ end of reporter transcripts, which can have significant effects on transcript stability and requires sensitive RNA ligation protocols. Fourth, these previous systems are universally encoded on plasmids to increase signal, but this leads to issues when trying to understand promoter function. In bacteria, plasmids exist at variable copy number, which can both contribute to expression noise^28^ and saturate endogenous transcriptional cofactors^29^. Finally, libraries of synthesized oligos characterized in these assays contain considerable amounts of sequence errors, and in certain MPRAs imperfect oligos cannot be distinguished from perfect sequences^21,30^.

Here we present a new MPRA system for studying promoter function in *E. coli* that addresses all of these concerns. To do this, we combined and extended several previous efforts to develop a new high-efficiency, site-specific genomic recombination-mediated cassette exchange (RMCE) system capable of integrating large libraries of genetic designs into precise genomic locations. We then build a reporter system capable of exploring transcriptional activity using RNA-Seq while maintaining our ability to differentiate perfect from imperfect reporter sequences, similar to MPRAs in mammalian systems. We use this new MPRA system to integrate over 300,000 reporters and measure expression of 10,898 σ70 promoter variants, which are specifically designed to explore the relationships between the ‐10, ‐35, UP, and spacer elements across different sequence backgrounds. We show that this genomically-based reporter assay achieves robust, quantitative measurements of promoter strength and use these measurements to develop statistical models that predict promoter strength based on sequence element composition. Furthermore, we leverage the insight gained from these statistical models to identify and dissect higher order interactions between σ70 sequence elements.

## Results & Discussion

### Design and Testing of High-efficiency Genomic RMCE

We designed our MPRA system to be genomically encoded at a defined genomic locus while at the same time allowing for easy construction of large libraries. To this end, we first used lambda-red recombination to insert an engineered landing pad into six intergenic loci distributed at different distances and orientations from the origin of replication that were previously identified as potential landing pad insertion sites^31^ (Figure S1). This landing pad contains an engineered operon encoding both red fluorescence (mCherry) and chloramphenicol resistance (*cat*^*R*^). The operon is engineered to be exchanged in whole using the mutant loxP sites *loxm2/66* and *lox71*^32-34^, which have been previously shown to mediate high-efficiency exchange from the genome using the GETR system^32^. Briefly, these two sites allow for the subsequent cassette exchange with a vector containing complementary lox sites, *loxm2/71* and *lox66*. *lox66* and *lox71* sites are capable of undergoing Cre-mediated cassette exchange, and their recombination irreversibly produces the inactive lox site, *lox72.* Furthermore, the *m2* mutation alters the spacer sequence to make them incompatible with natural spacer sequences, thereby preventing cis-recombination events^34^. To direct RMCE to the landing pad, we designed and constructed an integration vector composed of an arabinose-inducible Cre recombinase^35^, a temperature-sensitive origin-of-replication^36^, and a modular payload flanked by *loxm2/71* and *lox66* sites complementary to those in the landing pad (Figure S1). We can replace the landing pad cassette with our engineered design by transforming the integration vector into a strain engineered with the landing pad, inducing Cre-mediated recombination, and selecting for integrated cells while simultaneously removing unintegrated plasmid with heat-curing. We used flow cytometry to track integration of an sfGFP fluorescent marker into the *nth-ydgR* locus at high efficiency (Figure 1A). Initially, cells only express mCherry, but upon introduction of a constitutive donor plasmid as a library, and induction of Cre recombinase, we find that almost two-thirds of the cellular population undergo cassette exchange. Since the donor cassette also includes a resistance marker, when we subsequently apply selection 94.3% of the population contained the reporter and had lost expression of genomic mCherry, indicating proper cassette exchange.

To evaluate potential location-specific effects on expression, we measured mCherry expression from different integrated loci within the *E. coli* genome using flow cytometry (Figure 1B). We used six previously described locations with two flanking the origin of replication (*atpI-gidB, yieN-trkD)*, two located midreplichore (*ybbD-ylbG, essQ-cspB*), and two near the terminus (*nth-ydgR*, ygcE-ygcF)^31^. All six pairs are located on opposite sides of the genome, and face the direction of DNA replication. We found that the two sites near the origin varied greatly, with the *yieN-trkD* locus having 426% higher median expression than the *atpI-gidB* locus (Figure 1B). This may be due to differences in copy number around the origin^37^ or collisions with replication machinery^38^. The pairs of mid-replichore and termini loci only varied by 4.2% and 4.6% between each other respectively, and the median expression was approximately 27% higher at the termini location as compared to the mid-replichore locations. Finally, we tested expression of landing pads engineered in both orientations at the *nth-ydgR* locus and observed little difference in expression as has been previously observed^37,39^ (Figure 1C).

**Figure 1.**
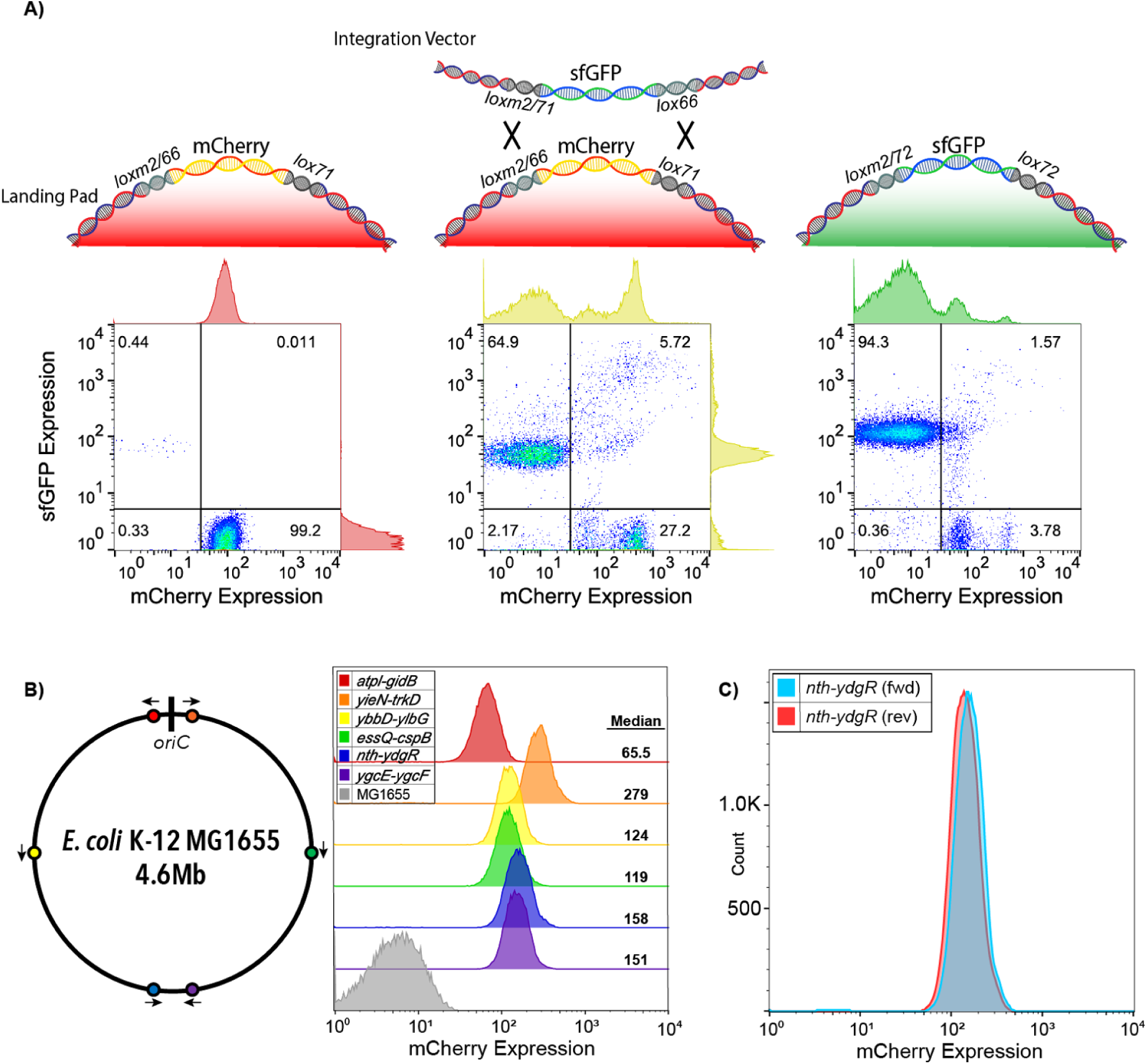
Recombination-mediated cassette exchange (RMCE) allows for high-efficiency genomic integration. A) We developed a cre-lox based RMCE that utilizes a combination of asymmetric (*lox66 and lox71*) and incompatible (*lox and loxm2*) loxP sites to allow for RMCE. We tracked the cell population with flow cytometry during RMCE. *Left*: Population of cells containing the mCherry landing pad engineered in the *nth-ydgR* locus prior to RMCE. *Center.* After transformation and RMCE of constitutively expressed sfGFP library, but prior to selection, both exchanged and unexchanged populations co-exist showing that an estimated two-thirds of the cells undergo RMCE. *Right:* Post-selection population shows 94.3% of the resultant population contains the cassette (as measured by constitutive sfGFP expression) and loss of the original landing pad mCherry expression. B) Expression of mCherry landing pads at six previously characterized locations spanning the *E. coli* genome^31^. Arrows indicate the landing pad orientation. C) Comparison of mCherry expression from the landing pad in both orientations at the *nth-ydgr* locus.

### Reporter and Library Design, Construction and Testing

To test the transcriptional output of large libraries of reporters in multiplex we designed and built a reporter construct. The final RMCE donor cassette (Figure S1) contains the promoter to be tested, a RiboJ self-cleaving ribozyme sequence^40^, and an sfGFP reporter with a random 20nt barcode in the 3’ UTR that uniquely identifies the promoter variant followed by a transcriptional terminator^41^. The RiboJ sequence standardizes the 5’ UTR of the reporter, decoupling transcriptional activity from any potential stability effects different UTR regions might have^40^. Immediately downstream, a constitutive promoter drives expression of the kanamycin resistance gene (*kan*^*R*^), allowing for selection of the RMCE donor cassette. The entire RMCE donor cassette is flanked by transcriptional terminators that isolate the reporter from local transcription events that might occur outside the reporter cassette. The barcodes are constructed by first amplifying the promoter library with a primer that adds a random 20nt barcode downstream. We subsequently clone the library of barcoded promoters into the RMCE donor plasmid and we use paired-end, next-generation sequencing to map the relationships between barcodes and variants. This approach allows us to identify promoters that contain sequence errors so that we may filter them out of downstream analyses. Finally, we clone the constant RiboJ::sfGFP sequence between the promoter and barcode. We engineered several other aspects of this cassette including restriction-enzyme sites for high-efficiency cloning and a priming site downstream of the barcode to facilitate reverse transcription and sequencing of the barcode.

We designed a library of 12,288 σ70 promoters to explore every possible combination of a set of 3 UP elements, 8 ‐35 regions, 8 spacer sequences, 8 ‐10 regions, and 8 background sequences (Figure 2A). The UP^13^, ‐35, and ‐10 elements^42^ selected span a large range of previously characterized activities. We chose to study these elements because they physically anchor RNAP to the promoter^43^ and we were interested in deconstructing how they they individually and cooperatively contribute to expression through this mechanism and others. We designed the spacer sequences to have variable GC content and flexibility, which have been shown to influence promoter expression^10^. Due to the position of the spacer between the ‐35 and ‐10 elements, we hoped to capture unique effects between spacer variants and core promoters of variable quality. Lastly, we extracted the backgrounds from non-promoter, intergenic regions of the *E. coli* genome that vary in GC content. Although sequences within the promoter background have been previously shown to modulate expression^44-48^, we varied this sequence primarily as a control to see how consistent observations were between sequence contexts. In addition to our library of σ70 promoter variants, we included 470 negative controls, which are intergenic regions that appear to be transcriptionally quiescent in RNA-Seq studies^49-51^.This library was synthesized, cloned, and integrated the library using this RMCE method into the *nth-ydgR* locus in the same direction as the DNA replication (Figure 2B). We chose this locus due to its proximity to the replication terminus which is present at a lower copy-number in rapidly dividing *E. coli*^52,53^. During the barcode mapping stage, we found 351,275 unique promoter-barcode combinations. After RMCE, we detected 318,825 (90.5%) of those barcodes using RNA and DNA-Seq. We also did not see large distortions in the overall distribution of barcodes per variant or the number of DNA reads per barcode before and after integration (Figure 2C, S2).

**Figure 2.**
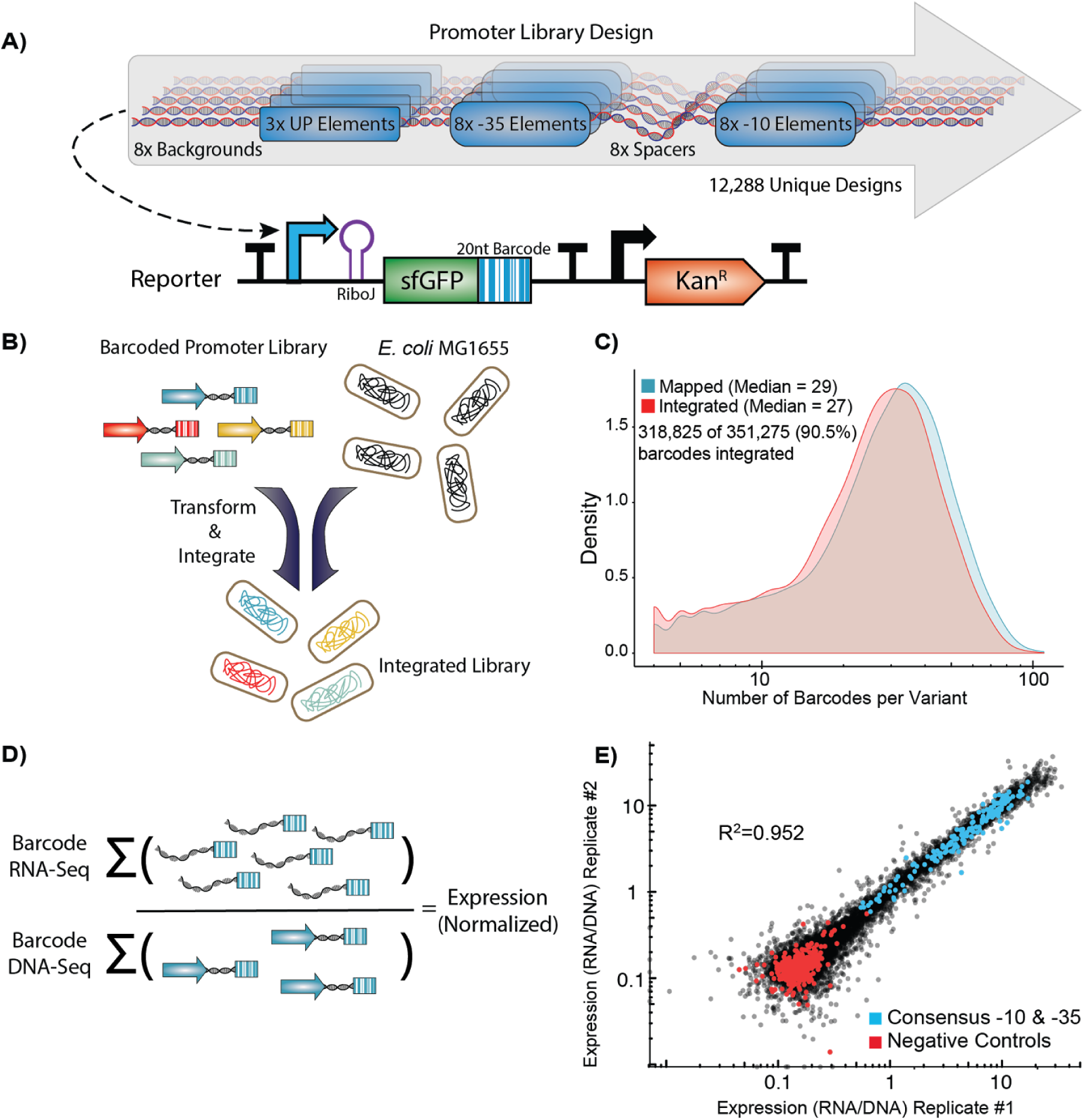
High-throughput quantification of σ70 promoter strength. A) We designed and constructed a σ70 promoter library using an oligonucleotide microarray, and cloned the library into a custom-made reporter construct. The reporter contains a promoter to be tested, a RiboJ self-cleaving ribozyme sequence to standardize the reporter 5’ UTR, and an sfGFP coding sequence followed by a 20 nt barcode in the 3’ UTR that identifies the promoter variant. The exchange cassette also includes a constitutive kanamycin resistance marker downstream of the reporter for selection purposes. B) Pooled promoters are uniquely barcoded using PCR, cloned into the exchange vector, and integrated into the *E. coli nth-ydgR* locus as a library. C) Pre-integration barcodes are identified during mapping stage and integrated barcodes are identified when quantifying promoter strength using RNA-Seq and DNAseq. We found 90.5% of the barcodes that were observed in the mapping stage (blue histogram), were later observed in the integrated library (red histogram), and the overall distributions remained similar. D) Expression of each promoter is calculated as the sum of all RNA counts divided by the sum of all DNA counts for all barcodes mapped to a given promoter. E) Promoter strength measurements are highly correlated (*R*^2^=0.952, p < 2.2×10^-16^) between technical replicates and discriminate between negative controls and promoters with consensus core elements.

To measure the expression of each promoter, we grew the library to exponential phase in defined media before extracting and sequencing both RNA and DNA barcodes. To account for differences in the abundance of each barcoded promoter, we calculated expression by normalizing the number of RNA counts to the number of DNA counts for all barcodes mapped to a single promoter (Figure 2D). In total, we performed three biological replicates in which the promoter library was grown on three occasions in separate cultures before being processed for RNA and DNA sequencing of the barcodes. In addition, we performed technical replicates of one sample in which two RNA and DNA samples from a single culture of the library were processed in parallel for sequencing. Expression of our promoter variants spanned a 100-fold range and were highly consistent between biological replicates (*R*^2^=0.952, *p* < 2.2 × 10^-16^) (Figure 2E). In addition, we observed a predictable segregation between our negative controls and promoter variants containing consensus ‐10 and ‐35 elements. We found a large spread in promoter activity even amongst those promoters containing consensus ‐10 and ‐35 sequences, with the strongest promoter containing consensus ‐10 and ‐35 regions having 29.9-fold higher activity than the weakest (Figure 3A). In general, for all data we see strong trends of closer to consensus ‐10 and ‐35 regions generally increasing transcription strength of the promoters (Figure 3B). However, there are many exceptions and the variance between promoters with identical UP, ‐10, and ‐35 regions can vary dramatically depending upon the the spacer and background sequences.

**Figure 3.**
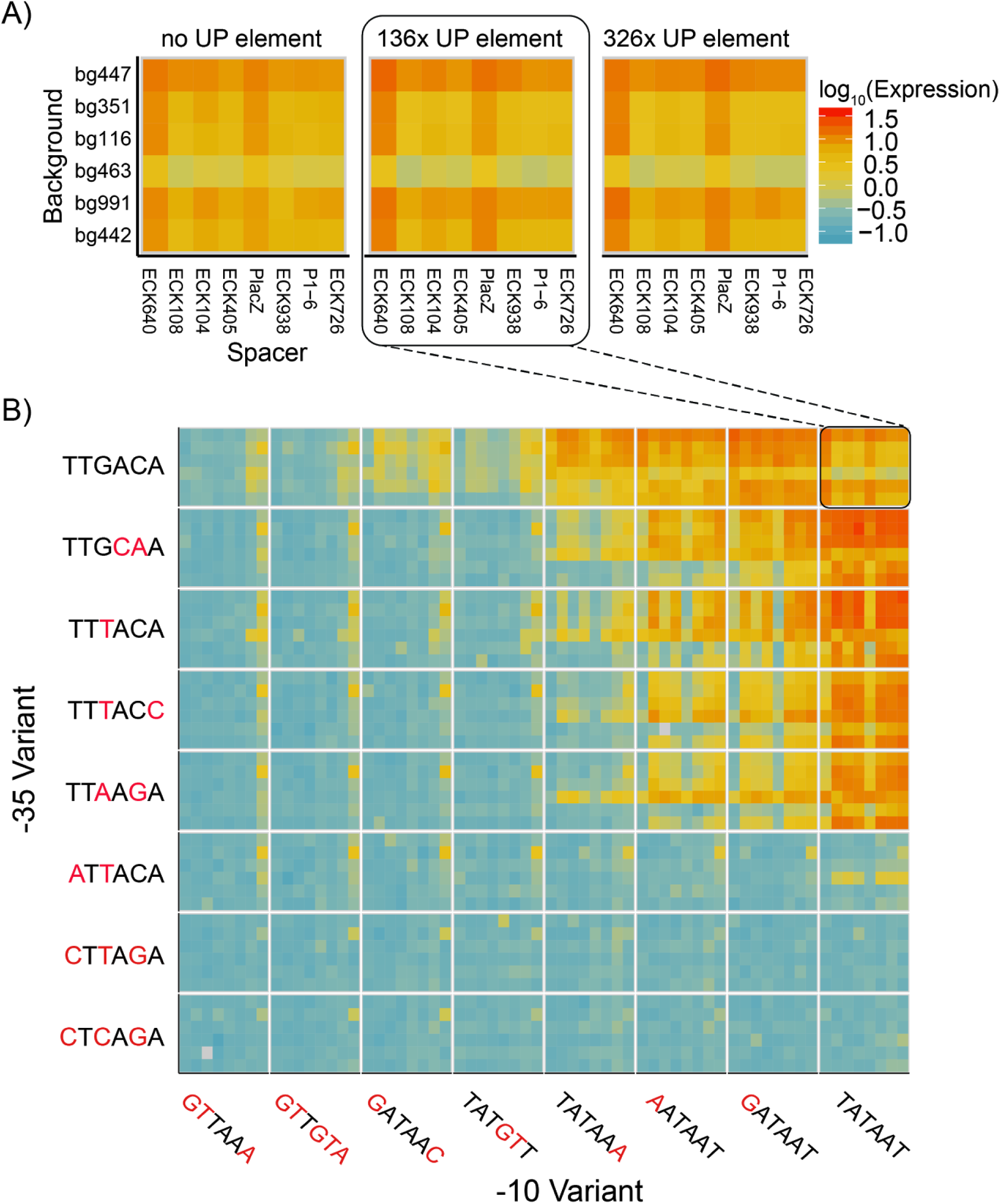
Expression levels for thousands of promoters. A) We plot the expression of all the promoters containing consensus ‐10 and ‐35 elements we measured in the library (*red* to *blue* is an estimated 100 fold decrease in measured expression). Each block of 48 squares displays six different backgrounds vertically against eight different spacer sequences horizontally. The three blocks represent the UP element choices used. We did not display two backgrounds for space and because they contained the most missing data but have included them in the supplement (Figure S3). The expression levels vary up to 29.9-fold based based on different background, spacer, and UP element choices. B) We plot expression of 3,072 promoters with the 136x UP element in blocks of 48 measurements (as in 3A), but now with all ‐10 (horizontal) and ‐35 (vertical) choices we measured in our assay. Expression generally increases as the ‐10 and ‐35 elements approach the consensus, yet like the consensus, there is variance amongst promoters with the same ‐10 and ‐35 elements. Promoter variants for which we could not detect more than four unique barcodes were omitted from our analysis and are displayed as grey squares.

### Promoter Activities can be Predicted by Sequence Element Combinations

Our approach provides us with a unique, large-scale training set of robust, quantitative measurements of promoter strength. A previously developed biophysical model of constitutive σ70 activity was modestly predictive of expression, (*R*^2^=0.351, p < 2.2 × 10^-16^) (Figure S4). Independent of thermodynamic prediction, we asked whether combinations of sequence elements, as we define them, can be used to predict expression by training statistical models. We first trained a multiple linear model on 50% of the promoter variants using the identities of the ‐10, ‐35, UP, Spacer, and Background as categorical variables and achieved an *R*^2^ = 0.395 (*p* < 2.2 × 10^-16^) on the remaining data (Figure S5A). Based on previous studies showing interaction between the ‐10 and ‐35 regions^14^, we included an interaction term between the ‐10 and ‐35 region, and *R*^2^ increased to 0.611 (*p* < 2.2 × 10^-16^) (Figure S5B). Previous studies of σ70 element interactions have shown that these elements primarily modulate expression in a multiplicative manner^43^, consistent with simple thermodynamic binding models of the ‐10 and ‐35 regions with RNA polymerase^42^. Therefore, we hypothesized a multiple linear regression on log-transformed data may adequately capture the relationship between promoter element composition and their resulting expression. Indeed, the log-linear statistical models worked better both with (*R*^2^ = 0.799, *p* < 2.2 × 10^-16^) and without the ‐10 and ‐35 interaction term (*R*^2^ = 0.596, *p* < 2.2 × 10^-16^) (Figures 4A & S5C). ANOVA revealed that a vast majority of expression variance in our dataset (73.7%) and the model’s power (90.9%) could be explained solely by the identity of the ‐10 and ‐35 elements and the interactions between them (Figure 4B). The remaining terms added only 7.4% of the variance explained, with approximately 19% of the variation in our dataset remaining unexplained. While this may indicate these other elements affect promoter activity very little, the overall dataset shows some patterns and the unexplained variance may be due to more complex cooperative relationships between elements.

**Figure 4.**
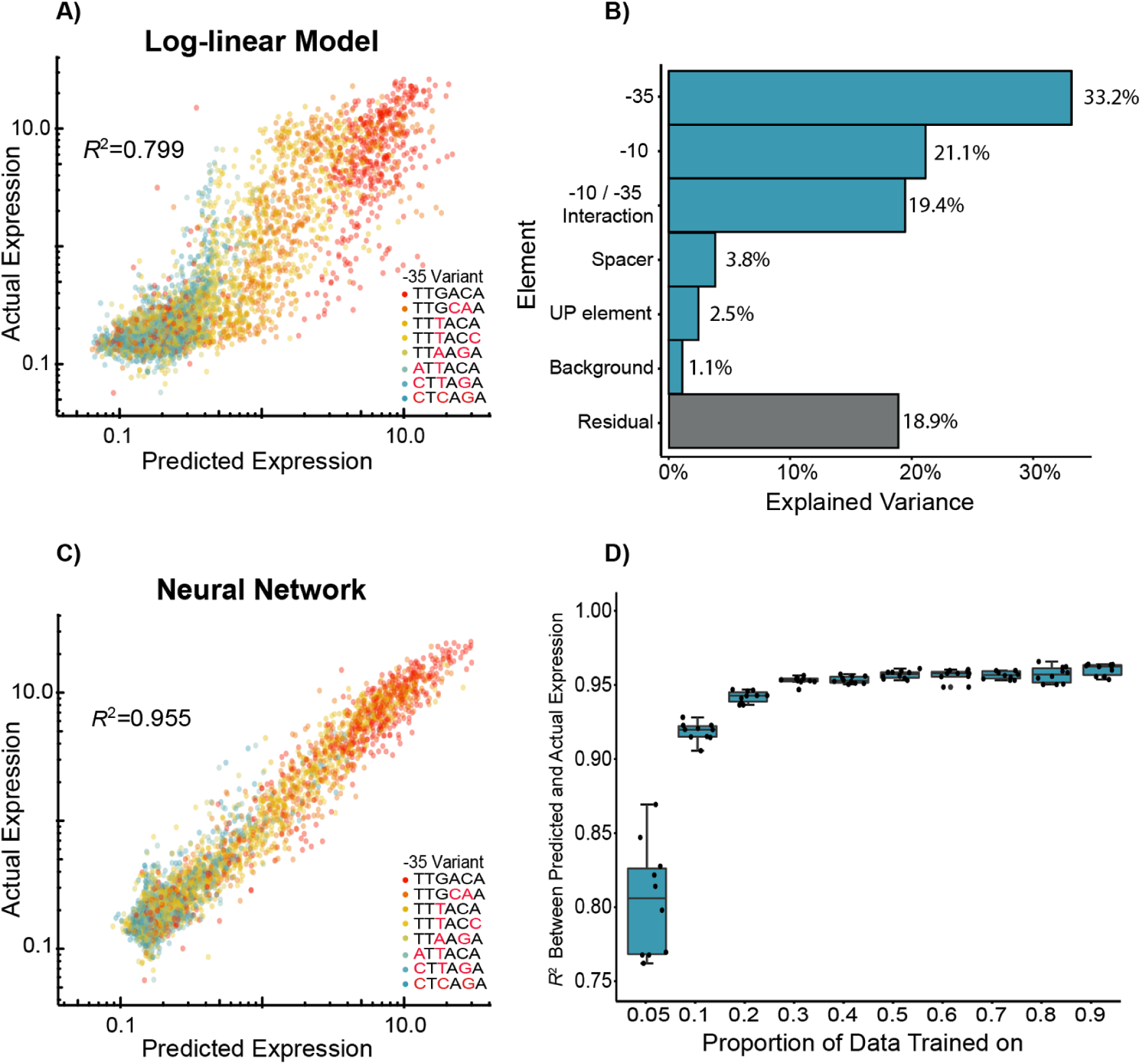
Predictive modeling of σ70 promoter strength. A) We trained a log-linear model on 50% of the data, and the resultant predictions on the remaining data explain approximately 80% of the variance in expression within our dataset. B) We analyzed the model by ANOVA and found that approximately 73.7% of variance in promoter expression can be explained by the ‐10 and ‐35 elements (and their interaction). C) We also trained a simple neural network model and found that the resultant predictions captured an estimated 95.5% of the promoter variance, indicating that these models are better able to capture more complex interactions between sequence elements. D) We trained the same neural network models with 10-fold cross-validation and show that we can effectively predict promoter expression when trained on as little as 5% of the data. In 4A, 4C, and 4D, *R*^2^ is the coefficient of determination between predicted and actual expression values on the held-out datasets.

To address the possibility for more complex nonlinear interactions, we implemented a simple neural network (NN) statistical model (Figure S6) with the hypothesis that these types of networks may pull out more subtle effects of element combinations. When trained on 50% of our data, the NN model was able to explain 95.5% (*p* < 2.2 × 10^-16^) of the variance in our dataset (Figure 4C). Surprisingly, by training on various proportions of our data we found that this neural network explained 94.2% of the variation when trained on 20% of the data (Figure 4D), and even training on 5% of the data produced NN models on par with our log-linear models (Figure S5D). The success of these NN models confirmed our hypothesis that while the ‐10 and ‐35 regions behave in a fairly predictable manner, the remaining unexplained variance is likely due to more complex relationships than can be easily captured by linear relations.

### Identifying Complex Interactions Between Sequence Elements

Because of the difficulty in interpreting neural networks, we examined specific combinations of elements to better understand these more complex relationships. Based on our results from the linear model, we analyzed the relationship between ‐10 and ‐35 variants. A clear trend emerged where the overall expression of the library increased as the ‐10 variant approached the consensus sequence (Figure 5A). Despite this trend, the most active promoters consisted of variants with consensus ‐35 sequences whose ‐10 sequence deviated from the consensus by a single nucleotide. Pairwise analysis of all ‐10 and ‐35 combinations confirmed that for both elements, expression was highest when one, but not both elements matched the consensus (Figures 3B and 5B).

**Figure 5.**
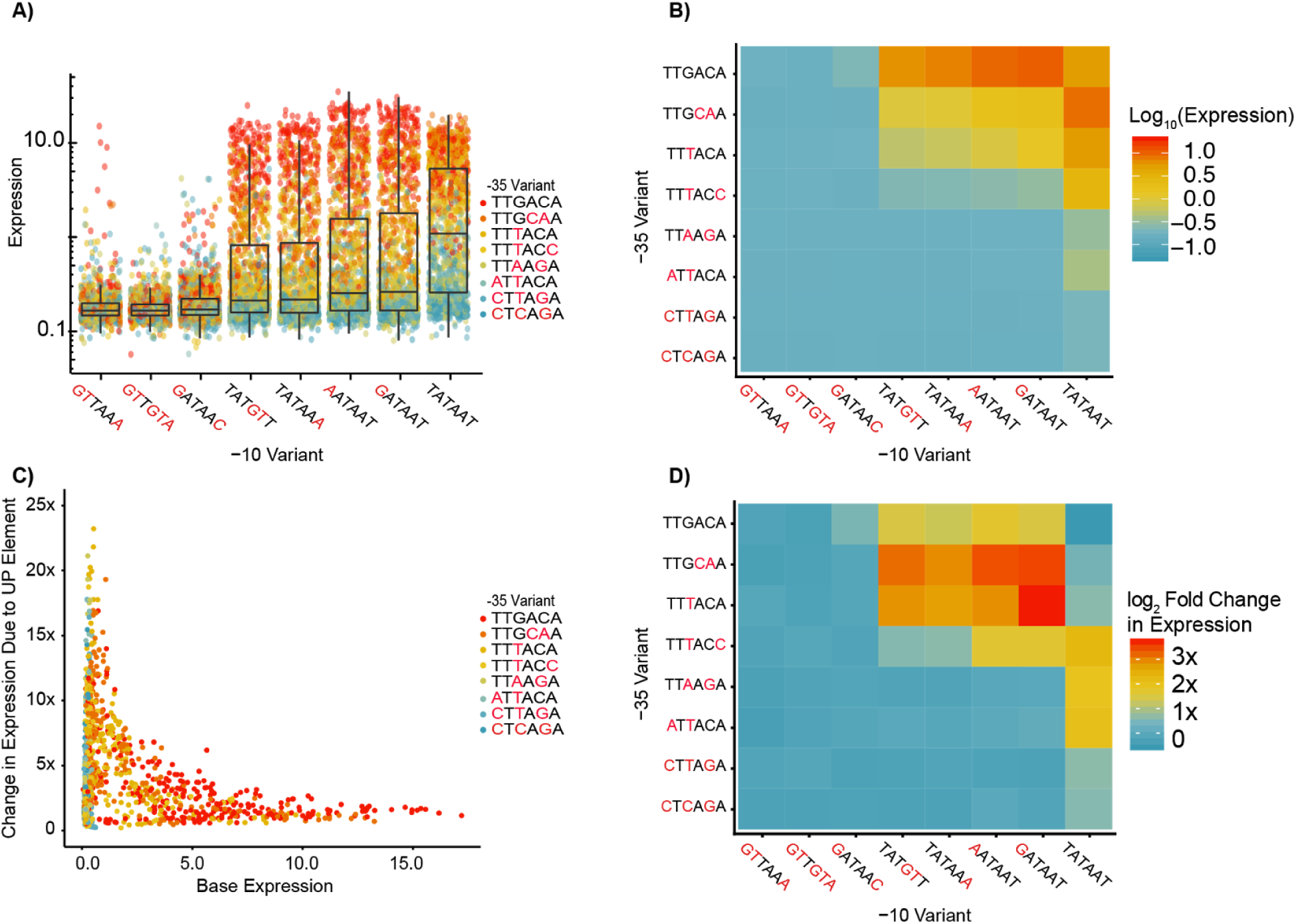
Identification of nonlinear interactions among promoter elements with direct RNAP Interactions. A) We plot all promoters split by ‐10 element and colored by ‐35 element. The overall promoter expression increases approaching the consensus ‐10 and ‐35, yet the strongest expressing promoters with a consensus ‐10 tend not to be those with a consensus ‐35. B) The median expression of all promoters as a function of the ‐10 and ‐35 identity shows a similar general trend towards increased expression as ‐10 and ‐35 gets closer to consensus. However, median expression of promoters containing a combination of a consensus and mutant ‐10 and ‐35 elements is higher than promoters containing both consensus sequences. C) We plot the fold-change increase in expression due to the addition of the 326x UP element as a function of the expression of the promoter without the UP element. Weaker promoters have the greatest increase in expression upon addition of the consensus UP element. D) We show the median log_2_ fold-change in expression for all ‐10 and ‐35 element combinations upon addition of the 326x UP element. On average, expression of promoters containing consensus ‐10 and ‐35 elements drops by 15%.

In addition to the ‐10 and ‐35 elements, the UP element is another point of physical contact between RNAP and the promoter^11,43^. Considering this, we postulated that addition of an UP element would serve a compensatory role for promoters with weak ‐10 and ‐35 elements. We observed a clear trend where weaker promoters received the greatest benefit upon addition of the consensus UP element (Figure 5C). In addition, we found several cases where promoters decreased in expression upon receiving the UP element, and a majority of these promoters contained the consensus ‐10 variant (Figure S7). It has been proposed that a strong UP element may decrease transcription of some promoters by inhibiting promoter escape^13,54^, which may explain our observation. Despite the clear trend we observed, the consensus UP element had highly variable effects when added to different combinations of ‐10 and ‐35 elements (Figure 5D). The weakest combinations of ‐10 and ‐35 elements did not receive the greatest increase in expression upon addition of the consensus UP element. Instead, the strongest non-consensus ‐10 and ‐35 elements had the greatest increase and this effect weakened as these core elements deviated further from the consensus. Also, addition of the consensus UP element enabled expression from promoters with otherwise inactive ‐35 variants, which has not been observed in the absence of an extended ‐10 motif (Figure S8)^43,54^.

The background sequences as we define them include many other regions known to affect transcription, including the discriminator and initial transcribed region^26,44-48^. We found modest differences in the distribution of expression for each background, and this appeared to be unrelated to background GC content (Figure 6A). However, expression of promoters with consensus ‐10 and ‐35 elements varied between background suggesting that there exists context-specific behavior amongst different compositions of core promoter elements. Further investigation revealed that many different combinations of ‐10 and ‐35 elements have variable expression between backgrounds, although the preferred promoters for each background are mostly consistent (Figure S9). To determine whether the background exhibited nonlinear interactions with promoter elements, we trained our neural network as before, but ignoring background. Although background sequence only accounted for 1.1% of the variance in our dataset according to the log-linear model, the performance of the neural network trained on 50% of the data dropped notably (*R*^2^ = 0.87, *p* < 2.2 × 10^-16^) (Figure S10). This suggests that nonlinear interactions with background sequences contribute a considerable amount to overall promoter expression.

**Figure 6.**
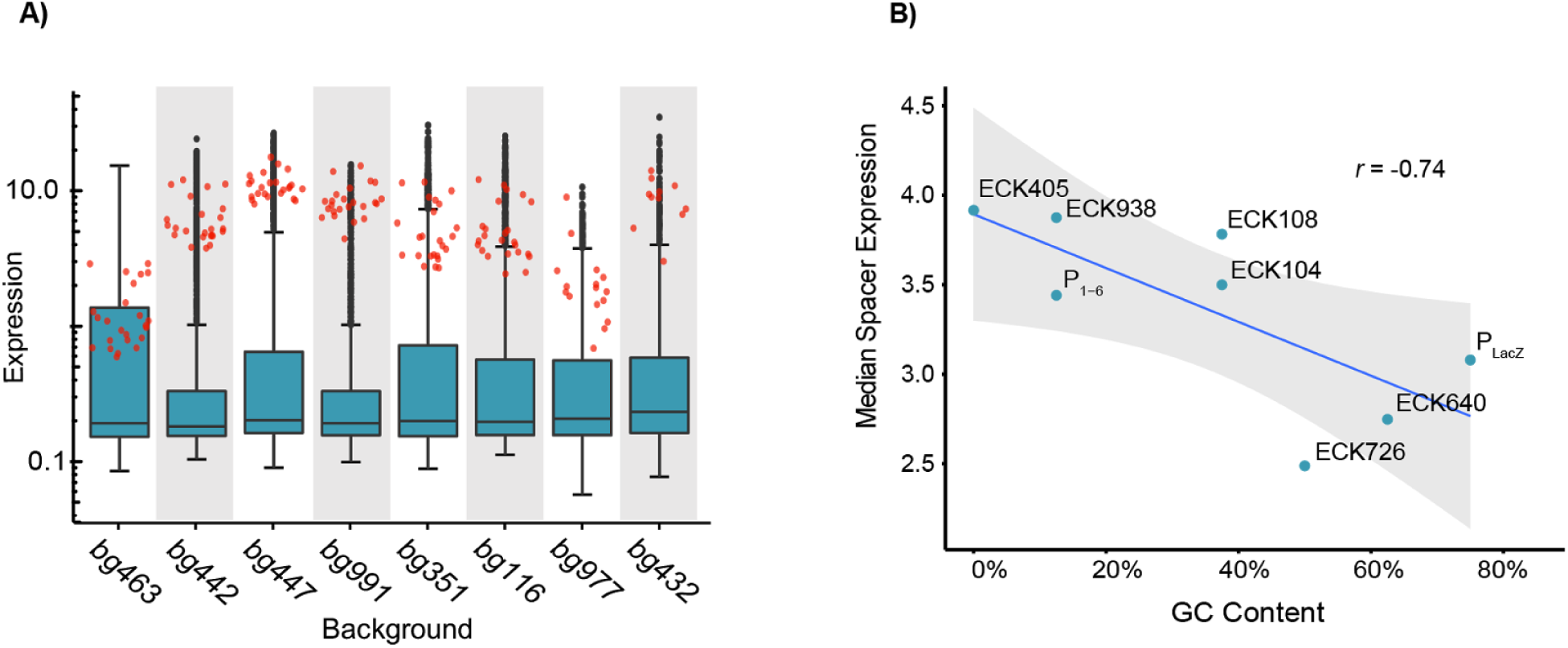
Effects of background and spacers on expression. A) The distribution of expression levels of promoters with different promoter backgrounds (*boxplots*) is similar yet consensus promoters (*red points*) vary drastically across these same contexts. Backgrounds are arranged from left to right by increasing GC content. B) The spacer GC content is negatively correlated with promoter expression. Each point represents the median expression amongst active promoters (RNA/DNA > 0.5) containing the indicated spacer. (*r* = ‐0.74, *p* =.036).

Finally, the spacer element partitions the ‐10 and ‐35 elements and has been suggested to contribute to expression through its GC nucleotide content^10^. We found a modest negative correlation between GC content in the ‐20 to ‐13 region of the spacer and promoter expression (*r* = ‐0.74, *p* = 0.036), though the effect size was small (Figure 6B). This may be due to reduced flexibility of the spacer inhibiting RNAP association or GC hydrogen-bonding impeding promoter melting^10,55,56^. One spacer variant, ECK726, had a unique effect in which it could stimulate transcription amongst promoters with otherwise unviable ‐10 and ‐35 combinations (Figure S11). Ultimately, we find evidence indicating that the ‐13 to ‐17 region of the spacer influences promoter activity and there are likely sequence-specific effects involving particular segments or nucleotides within the spacer.

## Conclusions

The full relationship between 150 nucleotide sequences used here and promoter activity can never be fully explored computationally, much less experimentally (4^150^ sequences). Building mechanistic and statistical models allow us to break down this complexity into separable shorter sequences that can be independently characterized and composed to predict function. The results of our combinatorial promoter library show that expression of σ70 promoters is primarily dictated by the identity of their ‐35 and ‐10 elements, which taken together are only 12 nucleotides in length. We also show that substantial nonlinear relationships arise from interactions with other sequence elements that can drastically alter expression. These nonlinear interactions can be captured by simple neural network models trained only on small proportions of the dataset, indicating that their effects can be accounted for through future study. However, it is likely that our use of only a few sequences for each of the elements underestimates their full complexity. Future work will focus on parameterizing how different sequences affect the strength of individual elements, quantifying the relationships between elements, and understanding the mechanistic basis for these patterns. Through this exploration, we hope to build predictive algorithms for promoter function that will be both useful for engineering purposes and the analysis of promoter function and evolution in microbial populations.

The platform we developed here enables site-specific integration of large libraries of reporter constructs in *E. coli.* Though we integrated over 300,000 unique promoter barcode combinations, we expect the methods are easily scalable and compatible with interrogating millions of variants. Furthermore, this approach should work for any of the many model systems in which Cre-recombination is available^57-59^ and for a wide variety of reporter assay formats, especially those which require single variants per cell. While the methods we use to generate landing pads and perform the subsequent cassette exchange have been previously established, to our knowledge we are the first to use them in this manner. Currently the process of cloning, integrating, and measuring the expression of large promoter libraries can be completed at a casual pace in six weeks and is amenable to rapid iteration. This platform should serve as an efficient and versatile means to conduct multiplexed reporter assays in the future to deconstruct the complex relationships between sequence and function.

## Acknowledgements

We thank Suhua Feng of the UCLA Broad Stem Cell Research Center for technical assistance with high throughput sequencing, Reid C. Johnson for technical advice, and Nathan B. Lubock for feedback and computational guidance.

## Abbreviations

Luria Broth (LB), superfolder Green Fluorescent Protein (sfGFP), Untranslated Region (UTR), MOPS (3-(*N*-morpholino)propanesulfonic acid), Genome Editing via Targetrons and Recombinases (GETR)

## Supplementary Information

Figures S1-S12. Tables S1-S3.

## Methods

### Landing Pad Generation

Landing pad strains were constructed using lambda-RED recombination in previously identified safe loci^31,60^. The landing pad construct (Figure S1) was assembled and flanked with one of six pairs of homology arms, corresponding to each of the 6 landing pad locations. At the *nth-ydgR* locus, two sets of homology arms were used to generate landing pads in the forward and reverse orientation. The linear landing pad DNA was genomically engineered into *E. coli* MG1655 K.12 harboring pTKRED (Genbank GU327533) following the published protocol^31^, with the exception of chloramphenicol being used for selection of successful recombinants. The pTKRED plasmid was heat-cured from identified recombinants by growth at 42°C on LB plates with chloramphenicol (34 ug/mL).

### Plasmid construction

The integration vector, pLibacceptorV2 (Figure S1) (Addgene id: 106250), was designed to include three primary components:

1) A library cloning site containing a selectable marker and flanked by mutant loxP sites
2) 2)An arabinose-inducible Cre-Recombinase^35^
3) A heat-sensitive origin of replication^37^

The library cloning site was ordered from IDT as a G-Block. The arabinose-inducible Cre system was amplified from pARC8-Cre^35^ and the temperature-sensitive origin of replication (tsORI) was amplified from pTKRED^31,35^. Fragments were assembled using a SGI-DNA Gibson Assembly^®^ HiFi HC 1-Step MasterMix (#GA1100-4☓10M).

### Landing Pad Integration Demonstration Using Flow Cytometry

Landing pad integration in Figure 1A was demonstrated by integrating a constitutively expressed sfGFP into the *E. coli nth-ydgR* locus. First, a landing pad was engineered in the *nth-ydgR* locus of K12 MG1655 *E. coli* in the reverse orientation following the landing pad generation protocol detailed above. A constitutively expressed sfGFP was cloned using restriction ligation into the pLibacceptorV2 RMCE cassette and 100 ng of this plasmid was transformed into the aforementioned landing pad strain and grown overnight at 30 °C in 100 mL of LB + kanamycin (25 ug/mL). The following day, 200 million cells (estimated by OD) were inoculated in 200 mL LB + kanamycin (25 ug/ml) + .2% (g/mL) Arabinose and grown at 30°C for 24 hours. From the 24 hours induced culture, 400 million cells were inoculated into separate 80 mL of LB + kanamycin (25 ug/mL). Once culture was grown at 30 °C for 2 hours (temperature permissive) and the other was grown at 42 °C (temperature impermissive) for approximately 1.5 hours before both reached an OD_600_ ≈ .5. Similarly, 10 uL of the landing pad strain grown overnight was seeded into 1 mL of LB + chloramphenicol (34 ug/mL) and grown at 30 °C for 2 hours. After reaching OD_600_ ≈ .5, each culture was placed at 4 °C for 30 minutes before being diluted 1:100 in phosphate buffered saline pH 7.4 (Thermo Scientific #10010023) and analyzed by flow cytometry using a BIO-RAD S3 Cell Sorter.

### Landing Pad Location Effects on Expression

To evaluate whether landing pad choice affects expression of constitutive promoters, several landing pads (Figure S1) were engineered into six loci previously characterized by Kuhlman T. and Cox E.^31^. To characterize expression at each landing pads, strains were grown overnight in 1 mL LB + chloramphenicol (34 ug/mL) at 37°C. The following day, 10 uL of each culture was seeded into separate 1 mL LB + chloramphenicol (34 ug/mL) cultures and grown for 1.5 hours at 37 °C before reaching OD_600_ ≈ .5. Upon reaching OD_600_ ≈ .5, cells were placed at 4 °C for 30 minutes before being diluted 1:100 in 1 mL PBS and analyzed using flow cytometry.

### Minimal Library Design

We designed a library of 12,288 σ70 promoters to explore every possible combination of a set of 3 UP elements, 8 ‐35 regions, 8 spacer sequences, 8 ‐10 regions, and 8 background sequences. A complete list of sequences is listed in Table S1.

The UP elements were identified by a modification of the SELEX procedure to identify protein-binding sites^11^. Two elements were selected which increased transcription 136 and 326-fold *in vivo* relative to the natural *rrnb* P1 UP element. An additional null (zero length) UP element was used. We incorporated extra bases from the flanks of the background sequence to maintain constant length for the entire library. We used eight ‐35 regions and eight ‐10 regions that were previously designed to span a wide range of promoter activity for the *E. coli lac* promoter^42^. These motifs were designed based on “information footprints” detailing the contribution of each nucleotide at each position to promoter strength, learned from a library of approximately 200,000 *lac* promoters mutagenized in a 75 bp region containing the cAMP Receptor Protein and RNAP binding sites^18^.

Spacer sequences were designed to span a range of GC content and flexibility, which both have been shown to influence promoter expression. Flexibility was calculated based on trinucleotide parameters learned from nucleosome-binding data^61^. All spacers are 17 bp in length which is considered to be the optimal length for σ70-dependent promoters^10^.

Backgrounds were extracted from the non-promoter regions of the genome. We randomly selected eight 150 bp genomic regions that were at least 200 bp away from a transcription start site on either strand. In addition, 470 negative controls were included - intergenic regions that appear to be transcriptionally quiescent in RNA-Seq studies^49-51^.

We synthesized 120 bp of upstream promoter region, 1 bp for the transcription start site (TSS) and 29 bp of the initial transcribed region (ITR) downstream of the TSS. Previous work has studied the preferred starting nucleotide and location relative to the ‐10 region^62^. Based on this, we used an “A” at the TSS and required 5 bp between the end of the ‐10 region and the TSS. The ITR is taken from the 150 bp background sequence and not a specific promoter.

All combinations of the elements described above were synthesized, in addition to the negative controls, resulting in 12,288 σ70 promoters. Relative motif location was maintained for each sequence. An example promoter schematic has been included in the supplement (Figure S12). Several restriction enzyme and priming sites were added to the termini for library amplification and cloning. Any assembled promoter sequences containing these restriction enzymes were removed.

### Minimal Library Cloning

The oligonucleotide library was constructed by Twist Biosciences and delivered lyophilized as a 26 pmol pool. The library was resuspended in 100 uL of TE pH 8.0 and 1 uL was amplified for 12 cycles using GU72 and GU116 with NEB Q5 High-Fidelity 2x Master Mix (#M0492L). Unless otherwise stated, all amplifications were performed using this polymerase mixture. This product was then ran on a 2% TAE agarose gel and approximately 200 bp amplicons were extracted using a Zymoclean Gel DNA Recovery Kit (#D4008). For barcoding, 1 ng of this eluate was amplified for 10 cycles using primers GU72 and GU73. Following cleaning using a Zymo Clean and Concentrator Kit (#D40140), the library was digested using NEB’s SbfI-HF and XhoI.

The plasmid backbone, pLibacceptorV2 was digested using SbfI-HF and SalI-HF with the addition of rSAP (NEB #M0371S). The digested library was ligated into pLibacceptorV2 using T7 DNA Ligase (NEB #M0318S), cloned into 5-alpha Electrocompetent *E. coli* (NEB #C2989K), and plated on LB + kanamycin (25 ug/mL) yielding approximately 1.1 million colonies estimated by plating concomitant dilution plates. After allowing for 24 hours of growth on plates, the library was scraped and resuspended in LB, and then 800 million cells (based on OD_600_) were inoculated in 450 mL LB + kanamycin (25 ug/mL) overnight. Unless stated otherwise, all plasmids were isolated using a Qiagen Plasmid Plus Maxiprep Kit (#12963) and concentrated using a Promega Wizard SV Gel and PCR Clean-up System (#A9281).

In order to clone the RiboJ::sfGFP reporter construct, the library was digested using NEB’s BsaI-HF and NheI-HF with the addition of rSAP. The reporter construct was digested using NEB’s BsaI-HF and NcoI-HF. Similarly to the previous cloning step, the reporter was cloned into the library using T7 DNA Ligase, cloned into 5-alpha electrocompetent *E. coli*, and plated on LB + kanamycin (25 ug/mL), yielding 8 × 10^5^ colonies. The completed plasmid library was isolated as stated above.

### Barcode Mapping

After cloning the barcoded library into pLibacceptorV2, we used Next-Generation Sequencing (NGS) to map promoters to their respective barcodes. Sequencing libraries were prepared through subsequent PCR reactions in which the first step adds custom sequencing primer sites while the second step adds P5 and P7 illumina flow cell adapter sequences. To limit the formation of chimeric species during amplification, we limit PCR to the exponential amplification^63^ phase as determined by qPCR. Initially, the barcoded library was amplified for 12 cycles from 10 ng of isolated plasmid using primers GU79 and GU60. Following a DNA Clean-Up using a Zymo Clean and Concentrator Kit, a second PCR was performed to add flow cell adapters to the amplified library. This PCR used 1 ng of the previously amplified library and was for 8 cycles with primers GU70 and one of either GU82 or GU83 for separately indexing replicates. Samples were submitted to the UCLA Technology Center for Genomics and Bioinformatics for sequencing using a 2×150 bp NextSeq 500. Between both replicates, 55,873,216 reads were acquired and used to determine promoter-barcode associations.

We next used the sequencing data to computationally map each promoter variant to its corresponding barcodes. Demultiplexed reads were paired using Paired-End reAd mergeR (PEAR v0.9.1, default settings). Custom python code was used to identify reads corresponding to perfectly synthesized promoters and their respective barcodes. Briefly, this code searched the first 150 bp of each read for perfect matches to library variants. For reads with perfect matches, the last 20 bp of each read (the barcode) was extracted and a list was compiled mapping each barcode to the most frequently associated library variant. A single barcode appears many times in the sequencing data, and we took steps to ensure a barcode consistently mapped to the same variant. We required that all variants mapped to a single barcode be within an edit distance (Levenshtein distance) of 5 from one another (five single bp changes between the two sequences). We determined this number by bootstrapping a distribution of the edit distance between any two random sequences in our variant library, and setting the threshold to the first percentile (1%) of this bootstrapped distribution. Additionally, each barcode had to appear at least three times in order to be considered for downstream analysis, which we reasoned would eliminate barcodes which contained sequencing errors. Raw sequencing data and promoter-barcode associations have been made available on NCBI’s Gene Expression Omnibus (GEO Accession no. GSE108535).

### Library Integration

The isolated plasmid library was digested with SalI-HF and NheI-HF to eliminate incompletely cloned plasmid before transformation into electrocompetent MG1655 with a landing pad engineered in the *nth-ydgR* locus and plating on LB + kanamycin (25 ug/mL), resulting in 16 Million colonies. Colonies were resuspended in LB and 800 million cells were inoculated into 250 mL LB + kanamycin (25 ug/mL) and grown overnight. Several 2 mL frozen aliquots were made of this overnight culture.

The library was integrated into the *nth-ydgR* locus as follows. A frozen aliquot of MG1655 with a landing pad engineered in the reverse orientation at the nth:ydgR locus was transformed with the library and grown overnight in 200 mL LB + kanamycin (25 ug/mL). Following overnight growth, 400 million cells of this culture were seeded into 250 mL LB + kanamycin (25 ug/mL) + .2% arabinose (g/mL) and grown for 24 hours. After integration of the library, the plasmid backbone was removed through heat-curing. From the 24 hour induced culture, 800 million cells were inoculated into 80 mL of LB + kanamycin (25 ug/mL) and grown at 42 °C for approximately 1.5 hours before reaching an OD_600_ =.3. Upon reaching exponential growth, 200 million cells from this culture library were plated and grown for 16 hours at 42 °C. Heat-cured plates were scraped and resuspended in LB and 400 million cells were inoculated into 200 mL LB + kanamycin (25 ug/mL). This culture, consisting of our integrated and heat-cured library, was grown overnight at 37 °C and several frozen 2 mL aliquots were made.

### Library Growth and Sequencing Library Preparation

For each biological replicate, A 2 mL frozen aliquot of the library was inoculated in 200 mL MOPS EZ-Rich Media (TEKNOVA #M2105) with .2% glucose (g/mL) and 25 ug/mL of kanamycin and grown at 30 °C overnight. The overnight culture was used to seed a new culture at OD_600_ = .0005 and grown for approximately 5.5 hours at 37 °C to an OD_600_ = .5. The culture was rapidly cooled to 0 °C in an ice slurry for two minutes. Three 50 mL aliquots were pelleted at 4 °C by centrifugation at 13,000xg for two minutes and the supernatant was poured out before snap-freezing the pellets in liquid nitrogen. Three 5 mL aliquots were prepared using the same approach.

### RNA and DNA library preparation

RNA was extracted from 50 mL library pellets using a Qiagen RNEasy Midi kit (#75142) and 45 ug of each extract was concentrated using a Qiagen Minelute Cleanup Kit (#74204). Barcoded cDNA was generated from 25 ug of each concentrated RNA extract using Thermo Fisher SuperScript IV (#18090010) primed with GU101. The manufacturer’s protocol was followed aside from extending the reaction time to 1 hour at 52 °C. The cDNA reaction was cleaned using a Zymo Research DNA Clean and Concentrator kit (#D40140) before amplification. Barcoded cDNA was amplified via PCR for 13 cycles using primers GU59 and GU102. This reaction was cleaned using a Zymo Research DNA Clean and Concentrator Kit and 1 ng of this reaction was used in a second PCR for indexing and addition of flow cell adapters. The second PCR was for 8 cycles and utilized primers GU102 and either GU61 or GU62.

gDNA was extracted from 5 mL cell library pellets using a Qiagen Gentra Puregene kit (#158567). Barcoded DNA was amplified from 1 ug of gDNA via PCR for 14 cycles using primers GU59 and GU60. The reaction was subsequently cleaned using a Zymo Research DNA Clean and Concentrator kit. To add sequencing adapters and indices to the library, 1 ng of this reaction was subject to a second PCR for 8 cycles using primers GU70 and either GU63 or GU64. RNA and DNA sequencing libraries were cleaned using a Zymo Research Clean and Concentrator Kit before quantification using an Agilent Tapestation.

In total, three biological replicates of the library RNA/DNA-seq were performed in which each replicate was separately grown to log phase before sequencing library preparation. For one biological replicate, two RNA/DNA extractions (technical replicates) were performed in parallel and sequenced together. The RNA and DNA of the other two biological replicates were sequenced altogether. All libraries were submitted to the Broad Stem Cell Research Center at UCLA for sequencing on a HiSeq2500. Raw sequencing data and promoter expression measurements have been made available on NCBI’s Gene Expression Omnibus (GEO Accession no. GSE108535). For each sample, the number of reads acquired are shown in Table S2.

### Barcode Measurement Processing

Barcode counts were extracted from demultiplexed replicate RNA and DNA reads using a custom bash script. From each sequencing file, the first 20 nucleotides (containing the barcode) were extracted, reverse complemented, and the counts for each unique sequence were determined. Each file was read into R studio (Version 1.0.153). Read counts were normalized using the following formula:

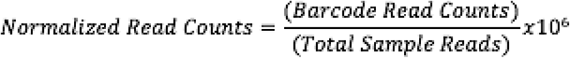

Normalized files were merged together based on common barcodes (`dplyr` package Version 0.7.2), generating a dataset containing normalized read counts for each barcode in each sample. This file was subsequently merged into the mapping file containing the list of barcodes and their mapped promoter (available on GEO Accession no. GSE108535 as file barcode_mapping.txt)

### Promoter Expression Quantification

Barcodes mapped to common promoters were aggregated and promoters that had fewer than four barcodes detected in either RNA or DNA sample amongst all replicates were removed from analysis. Of the 12,849 mapped promoters, 11,368 passed this threshold. To calculate promoter expression, promoter expression in each replicate RNA extraction was calculated as the sum of all RNA counts divided by the sum of all DNA counts for all barcodes mapped to that promoter.

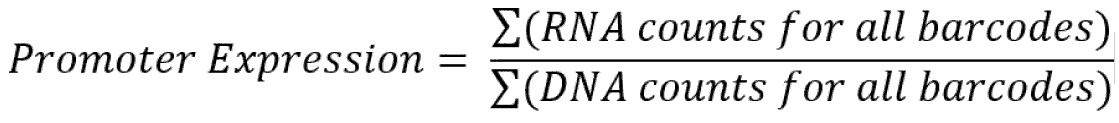

For the biological replicate in which technical replicates were performed, the mean expression of the technical replicates was calculated before averaging this biological replicate with the other two remaining biological replicates. This final average was used for all data analysis and modeling.

### Modeling

First, we randomly split our library of promoter variants into 50% training and 50% testing data sets. We fit the linear model using the `lm`() function in R (stats package version 3.3.3). We modeled expression based on the variant identity of each element and included an interaction term for the ‐10 and ‐35 elements. We used the `aov`() function (stats package) to calculate the model variance and the variance explained by each sequence element was calculated as the percentage of the sum of the squared deviation.

We used the R package `nnet` to fit our data to a single-hidden-layer neural network. The network topology was structured as 30 input nodes, each representing an element variant, 10 nodes in the hidden layer, and a single linear output node. The network was trained for 300 iterations with weight decay set to .01. We performed 10-fold cross-validation using the same network trained on different proportions of the training data, resampled between each fold and tested on the remaining proportion of the training data.

The neural network visual was created in R using `plot.nnet`() (RPackage:’NeuralNetTools’), updated to be compatible with neural networks generated by the ‘nnet’ R package. This update is courtesy of (https://beckmw.wordpress.com/tag/nnet/).

We modified a previously developed mechanistic model of promoter activation that considers the thermodynamic binding energy of RNAP to the *lac* promoter^18,42^. The RNAP model is specific to the *lac* promoter and scores positions ‐41 to - 1 (where 0 denotes the transcription start site). This model is summarized in an energy matrix, where each nucleotide at each position is an experimentally determined energy value. The binding energy of RNAP to a specific sequence is determined additively by the matrix values, where more positive values indicate less favorable binding. The matrix contains an 18bp spacer, while our minimal library contains a 17bp spacer. To accommodate this discrepancy, we do not score the spacer segment of our library and compute a partial RNAP binding energy score. R. Brewster et al. implemented a thermodynamic model based on the binding energy matrix developed by Kinney et al.^18^ which predicts expression based on the binding of RNAP. However, the different spacer length prevents us from fully implementing their thermodynamic model.

---

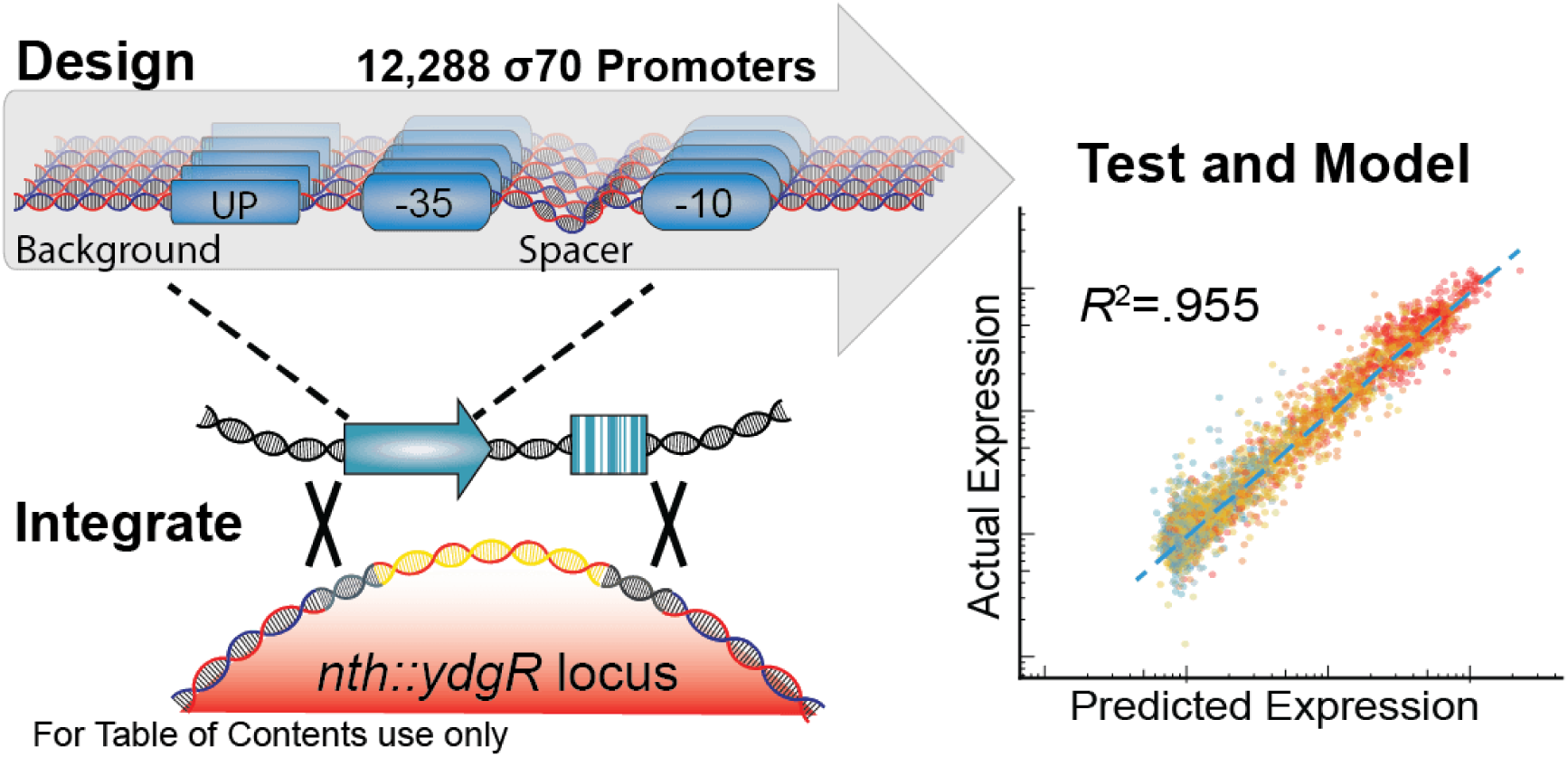

## References

(1) Berthoumieux, S., de Jong, H., Baptist, G., Pinel, C., Ranquet, C., Ropers, D., and Geiselmann, J. (2013) Shared control of gene expression in bacteria by transcription factors and global physiology of the cell. Mol. Syst. Biol. 9, 634.

(2) Browning, D. F., and Busby, S. J. W. (2016) Local and global regulation of transcription initiation in bacteria. Nat. Rev. Microbiol. 14, 638–650.

(3) Gerosa, L., Kochanowski, K., Heinemann, M., and Sauer, U. (2013) Dissecting specific and global transcriptional regulation of bacterial gene expression. Mol. Syst. Biol. 9, 658.

(4) Feklístov, A., Sharon, B. D., Darst, S. A., and Gross, C. A. (2014) Bacterial Sigma Factors: A Historical, Structural, and Genomic Perspective. Annu. Rev. Microbiol. 68, 357–376.

(5) Gruber, T. M., and Gross, C. A. (2003) Multiple sigma subunits and the partitioning of bacterial transcription space. Annu. Rev. Microbiol. 57, 441–466.

(6) Paget, M. S. B., and Helmann, J. D. (2003) The sigma70 family of sigma factors. Genome Biol. 4, 203.

(7) Jishage, M., and Ishihama, A. (1995) Regulation of RNA polymerase sigma subunit synthesis in Escherichia coli: intracellular levels of sigma 70 and sigma 38. J. Bacteriol. 177, 6832–6835.

(8) Jishage, M., Iwata, A., Ueda, S., and Ishihama, A. (1996) Regulation of RNA polymerase sigma subunit synthesis in Escherichia coli: intracellular levels of four species of sigma subunit under various growth conditions. J. Bacteriol. 178, 5447–5451.

(9) Campbell, E. A., Muzzin, O., Chlenov, M., Sun, J. L., Olson, C. A., Weinman, O., Trester-Zedlitz, M. L., and Darst, S. A. (2002) Structure of the Bacterial RNA Polymerase Promoter Specificity σ Subunit. Mol. Cell 9, 527–539.

(10) Liu, M., Tolstorukov, M., Zhurkin, V., Garges, S., and Adhya, S. (2004) A mutant spacer sequence between ‐35 and ‐10 elements makes the Plac promoter hyperactive and cAMP receptor protein-independent. Proc. Natl. Acad. Sci. U. S. A. 101, 6911–6916.

(11) Estrem, S. T., Gaal, T., Ross, W., and Gourse, R. L. (1998) Identification of an UP element consensus sequence for bacterial promoters. Proc. Natl. Acad. Sci. U. S. A. 95, 9761–9766.

(12) Carr, S. B., Beal, J., and Densmore, D. M. (2017) Reducing DNA context dependence in bacterial promoters. PLoS One 12, e0176013.

(13) Ross, W., Aiyar, S. E., Salomon, J., and Gourse, R. L. (1998) Escherichia coli promoters with UP elements of different strengths: modular structure of bacterial promoters. J. Bacteriol. 180, 5375–5383.

(14) Mutalik, V. K., Nonaka, G., Ades, S. E., Rhodius, V. A., and Gross, C. A. (2009) Promoter Strength Properties of the Complete Sigma E Regulon of Escherichia coli and Salmonella enterica. J. Bacteriol. 191, 7279–7287.

(15) Inoue, F., and Ahituv, N. (2015) Decoding enhancers using massively parallel reporter assays. Genomics 106, 159–164.

(16) White, M. A. (2015) Understanding how cis-regulatory function is encoded in DNA sequence using massively parallel reporter assays and designed sequences. Genomics 106, 165–170.

(17) Patwardhan, R. P., Lee, C., Litvin, O., Young, D. L., Pe’er, D., and Shendure, J. (2009) High-resolution analysis of DNA regulatory elements by synthetic saturation mutagenesis. Nat. Biotechnol. 27, 1173–1175.

(18) Kinney, J. B., Murugan, A., Callan, C. G., Jr, and Cox, E. C. (2010) Using deep sequencing to characterize the biophysical mechanism of a transcriptional regulatory sequence. Proc. Natl. Acad. Sci. U. S. A. 107, 9158–9163.

(19) Mogno, I., Myers, C. A., and Corbo, J. C. (2012) Complex effects of nucleotide variants in a mammalian cis-regulatory element. Proc. Natl. Acad. Sci. U. S. A. 109, 19498–19503.

(20) Melnikov, A., Murugan, A., Zhang, X., Tesileanu, T., Wang, L., Rogov, P., Feizi, S., Gnirke, A., Callan, C. G., Jr, Kinney, J. B., Kellis, M., Lander, E. S., and Mikkelsen, T. S. (2012) Systematic dissection and optimization of inducible enhancers in human cells using a massively parallel reporter assay. Nat. Biotechnol. 30, 271–277.

(21) Kosuri, S., Goodman, D. B., Cambray, G., Mutalik, V. K., Gao, Y., Arkin, A. P., Endy, D., and Church, G. M. (2013) Composability of regulatory sequences controlling transcription and translation in Escherichia coli. Proc. Natl. Acad. Sci. U. S. A. 110, 14024–14029.

(22) Goodman, D. B., Church, G. M., and Kosuri, S. (2013) Causes and Effects of N-Terminal Codon Bias in Bacterial Genes. Science 342, 475–479.

(23) Tewhey, R., Kotliar, D., Park, D. S., Liu, B., Winnicki, S., Reilly, S. K., Andersen, K. G., Mikkelsen, T. S., Lander, E. S., Schaffner, S. F., and Sabeti, P. C. (2016) Direct Identification of Hundreds of Expression-Modulating Variants using a Multiplexed Reporter Assay. Cell 165, 1519–1529.

(24) Vvedenskaya, I. O., Zhang, Y., Goldman, S. R., Valenti, A., Visone, V., Taylor, D. M., Ebright, R. H., and Nickels, B. E. (2015) Massively Systematic Transcript End Readout, “MASTER”: Transcription Start Site Selection, Transcriptional Slippage, and Transcript Yields. Mol. Cell 60, 953–965.

(25) Yus, E., Yang, J.-S., Sogues, A., and Serrano, L. (2017) A reporter system coupled with high-throughput sequencing unveils key bacterial transcription and translation determinants. Nat. Commun. 8, 368.

(26) Winkelman, J. T., Vvedenskaya, I. O., Zhang, Y., Zhang, Y., Bird, J. G., Taylor, D. M., Gourse, R. L., Ebright, R. H., and Nickels, B. E. (2016) Multiplexed protein-DNA cross-linking: Scrunching in transcription start site selection. Science 351, 1090–1093.

(27) Vvedenskaya, I. O., Vahedian-Movahed, H., Zhang, Y., Taylor, D. M., Ebright, R. H., and Nickels, B. E. (2016) Interactions between RNA polymerase and the core recognition element are a determinant of transcription start site selection. Proc. Natl. Acad. Sci. U. S. A. 113, E2899–905.

(28) Jahn, M., Vorpahl, C., Hübschmann, T., Harms, H., and Müller, S. (2016) Copy number variability of expression plasmids determined by cell sorting and Droplet Digital PCR. Microb. Cell Fact. 15, 211.

(29) Brewster, R. C., Weinert, F. M., Garcia, H. G., Song, D., Rydenfelt, M., and Phillips, R. (2014) The transcription factor titration effect dictates level of gene expression. Cell 156, 1312–1323.

(30) Kosuri, S., and Church, G. M. (2014) Large-scale de novo DNA synthesis: technologies and applications. Nat. Methods 11, 499–507.

(31) Kuhlman, T. E., and Cox, E. C. (2010) Site-specific chromosomal integration of large synthetic constructs. Nucleic Acids Res. 38, e92–e92.

(32) Enyeart, P. J., Chirieleison, S. M., Dao, M. N., Perutka, J., Quandt, E. M., Yao, J., Whitt, J. T., Keatinge-Clay, A. T., Lambowitz, A. M., and Ellington, A. D. (2013) Generalized bacterial genome editing using mobile group II introns and Cre-lox. Mol. Syst. Biol. 9, 685.

(33) Albert, H., Dale, E. C., Lee, E., and Ow, D. W. (1995) Site-specific integration of DNA into wild-type and mutant lox sites placed in the plant genome. Plant J. 7, 649–659.

(34) Langer, S. J., Ghafoori, A. P., Byrd, M., and Leinwand, L. (2002) A genetic screen identifies novel non-compatible loxP sites. Nucleic Acids Res. 30, 3067–3077.

(35) Eroshenko, N., and Church, G. M. (2013) Mutants of Cre recombinase with improved accuracy. Nat. Commun. 4, 2509.

(36) Hamilton, C. M., Aldea, M., Washburn, B. K., Babitzke, P., and Kushner, S. R. (1989) New method for generating deletions and gene replacements in Escherichia coli. J. Bacteriol. 171, 4617–4622.

(37) Block, D. H. S., Hussein, R., Liang, L. W., and Lim, H. N. (2012) Regulatory consequences of gene translocation in bacteria. Nucleic Acids Res. 40, 8979–8992.

(38) French, S. (1992) Consequences of replication fork movement through transcription units in vivo. Science 258, 1362–1365.

(39) Beckwith, J. R., Signer, E. R., and Epstein, W. (1966) Transposition of the Lac Region of E. coli. Cold Spring Harb. Symp. Quant. Biol. 31, 393–401.

(40) Lou, C., Stanton, B., Chen, Y.-J., Munsky, B., and Voigt, C. A. (2012) Ribozyme-based insulator parts buffer synthetic circuits from genetic context. Nat. Biotechnol. 30, 1137–1142.

(41) Chen, Y.-J., Liu, P., Nielsen, A. A. K., Brophy, J. A. N., Clancy, K., Peterson, T., and Voigt, C. A. (2013) Characterization of 582 natural and synthetic terminators and quantification of their design constraints. Nat. Methods 10, 659–664.

(42) Brewster, R. C., Jones, D. L., and Phillips, R. (2012) Tuning promoter strength through RNA polymerase binding site design in Escherichia coli. PLoS Comput. Biol. 8, e1002811.

(43) Hook-Barnard, I. G., and Hinton, D. M. (2007) Transcription initiation by mix and match elements: flexibility for polymerase binding to bacterial promoters. Gene Regul. Syst. Bio. 1, 275–293.

(44) Skancke, J., Bar, N., Kuiper, M., and Hsu, L. M. (2015) Sequence-Dependent Promoter Escape Efficiency Is Strongly Influenced by Bias for the Pretranslocated State during Initial Transcription. Biochemistry 54, 4267–4275.

(45) Henderson, K. L., Felth, L. C., Molzahn, C. M., Shkel, I., Wang, S., Chhabra, M., Ruff, E. F., Bieter, L., Kraft, J. E., and Record, M. T., Jr. (2017) Mechanism of transcription initiation and promoter escape by E. coli RNA polymerase. Proc. Natl. Acad. Sci. U. S. A. 114, E3032–E3040.

(46) Xue, X.-C., Liu, F., and Ou-Yang, Z.-C. (2008) A kinetic model of transcription initiation by RNA polymerase. J. Mol. Biol. 378, 520–529.

(47) Djordjevic, M., and Bundschuh, R. (2008) Formation of the Open Complex by Bacterial RNA Polymerase—A Quantitative Model. Biophys. J. 94, 4233–4248.

(48) Winkelman, J. T., and Gourse, R. L. (2017) Open complex DNA scrunching: A key to transcription start site selection and promoter escape. Bioessays 39.

(49) Thomason, M. K., Bischler, T., Eisenbart, S. K., Förstner, K. U., Zhang, A., Herbig, A., Nieselt, K., Sharma, C. M., and Storz, G. (2015) Global transcriptional start site mapping using differential RNA sequencing reveals novel antisense RNAs in Escherichia coli. J. Bacteriol. 197, 18–28.

(50) Conway, T., Creecy, J. P., Maddox, S. M., Grissom, J. E., Conkle, T. L., Shadid, T. M., Teramoto, J., San Miguel, P., Shimada, T., Ishihama, A., Mori, H., and Wanner, B. L. (2014) Unprecedented high-resolution view of bacterial operon architecture revealed by RNA sequencing. MBio 5, e01442–14.

(51) Gama-Castro, S., Salgado, H., Santos-Zavaleta, A., Ledezma-Tejeida, D., Muñiz-Rascado, L., García-Sotelo, J. S., Alquicira-Hernández, K., Martínez-Flores, I., Pannier, L., Castro-Mondragón, J. A., Medina-Rivera, A., Solano-Lira, H., Bonavides-Martínez, C., Pérez-Rueda, E., Alquicira-Hernández, S., Porrón-Sotelo, L., López-Fuentes, A., Hernández-Koutoucheva, A., Del Moral-Chávez, V., Rinaldi, F., and Collado-Vides, J. (2016) RegulonDB version 9.0: high-level integration of gene regulation, coexpression, motif clustering and beyond. Nucleic Acids Res. 44, D133–43.

(52) Slager, J., and Veening, J.-W. (2016) Hard-Wired Control of Bacterial Processes by Chromosomal Gene Location. Trends Microbiol. 24, 788–800.

(53) Fossum, S., Crooke, E., and Skarstad, K. (2007) Organization of sister origins and replisomes during multifork DNA replication in Escherichia coli. EMBO J. 26, 4514–4522.

(54) Ellinger, T., Behnke, D., Knaus, R., Bujard, H., and Gralla, J. D. (1994) Context-dependent Effects of Upstream A-Tracts. J. Mol. Biol. 239, 466–475.

(55) Hook-Barnard, I. G., and Hinton, D. M. (2009) The promoter spacer influences transcription initiation via σ70 region 1.1 of Escherichia coli RNA polymerase. Proc. Natl. Acad. Sci. U. S. A. 106, 737–742.

(56) Khandelwal, G., and Bhyravabhotla, J. (2010) A phenomenological model for predicting melting temperatures of DNA sequences. PLoS One 5, e12433.

(57) Araki, K., Araki, M., and Yamamura, K.-I. (2002) Site-directed integration of the cre gene mediated by Cre recombinase using a combination of mutant lox sites. Nucleic Acids Res. 30, e103.

(58) Trigg, S. A., Garza, R. M., MacWilliams, A., Nery, J. R., Bartlett, A., Castanon, R., Goubil, A., Feeney, J., O’Malley, R., Huang, S.-S. C., Zhang, Z. Z., Galli, M., and Ecker, J. R. (2017) CrY2H-seq: a massively multiplexed assay for deep-coverage interactome mapping. Nat. Methods 14, 819–825.

(59) Oberstein, A., Pare, A., Kaplan, L., and Small, S. (2005) Site-specific transgenesis by Cre-mediated recombination in Drosophila. Nat. Methods 2, 583–585.

(60) Yu, D., Ellis, H. M., Lee, E. C., Jenkins, N. A., Copeland, N. G., and Court, D. L. (2000) An efficient recombination system for chromosome engineering in Escherichia coli. Proc. Natl. Acad. Sci. U. S. A. 97, 5978–5983.

(61) Brukner, I., Sánchez, R., Suck, D., and Pongor, S. (1995) Trinucleotide Models for DNA Bending Propensity: Comparison of Models Based on DNaseI Digestion and Nucleosome Packaging Data. J. Biomol. Struct. Dyn. 13, 309–317.

(62) Liu, J., and Turnbough, C. L., Jr. (1994) Effects of transcriptional start site sequence and position on nucleotide-sensitive selection of alternative start sites at the pyrC promoter in Escherichia coli. J. Bacteriol. 176, 2938–2945.

(63) Thompson, J. R., Marcelino, L. A., and Polz, M. F. (2002) Heteroduplexes in mixed-template amplifications: formation, consequence and elimination by “reconditioning PCR.” Nucleic Acids Res. 30, 2083–2088.

